# The effect of liver enzymes on adiposity: a Mendelian randomization study

**DOI:** 10.1101/404319

**Authors:** Jun Xi Liu, Shiu Lun Au Yeung, Man Ki Kwok, June Yue Yan Leung, Shi Lin Lin, Lai Ling Hui, Gabriel Matthew Leung, C. Mary Schooling

## Abstract

Poorer liver function is positively associated with diabetes in Mendelian randomization (MR) studies. Observationally, adiposity is associated with poorer liver function. To clarify the etiology, we assessed the association of liver function with adiposity observationally and using two sample MR for validation.

In the “Children of 1997” birth cohort, we used multivariable linear regression to assess the associations of ALT and alkaline phosphatase (ALP) (IU/L) at ∼17.5 years with body mass index (BMI) (kg/m^2^). Using MR, genetic variants predicting ALT, ALP and gamma glutamyltransferase (GGT) (100% change in concentration), were applied to genome-wide association studies of BMI, waist circumference (WC) and waist-hip ratio (WHR) (standard deviations) to obtain unconfounded inverse-variance weighting estimates.

Observationally, ALT was positively associated with BMI (0.10, 95% confidence interval (CI) 0.09 to 0.11). ALP was inversely associated with BMI (−0.018, 95% CI −0.024 to −0.012). Using MR, ALT was inversely associated with BMI (−0.14, 95% CI −0.20 to −0.07), but not WC or WHR. ALP and GGT were unrelated to adiposity.

Poorer liver function might not cause adiposity; instead higher ALT might reduce BMI. Whether ALT contributes to diabetes by reducing muscle mass, given the no association of ALT with WC or WHR, requires investigation.

**Abbreviations:**

ALT: alanine transaminase
ALP: alkaline phosphatase
GGT: gamma glutamyltransferase
NAFLD: nonalcoholic fatty liver disease
BMI: body mass index
WC: waist circumference
WHR: waist-hip ratio
T2DM: type 2 diabetes mellitus
SEP: socioeconomic position
MR: Mendelian randomizatio
SNP: single nucleotide polymorphisms
GWAS: genome-wide association study
GIANT: Genetic Investigation of ANthropometric Traits
GIANTUKB: 2018 GIANT and UK Biobank meta-analysis
SD: standard deviation
IVW: inverse variance weighting
WM: weighted median

## Introduction

Observationally, poorer liver function, particularly nonalcoholic fatty liver disease (NAFLD), is associated with higher risk of type 2 diabetes mellitus (T2DM),^1,2^ but these studies are difficult to interpret because of the difficulty of distinguishing between correlated measures of liver function and the possibility of confounding by poor health causing both poor liver function and T2DM.^3^ Recently, Mendelian randomization (MR) studies have clarified that higher alanine aminotransferase (ALT) rather than other measures of liver function, could play a role in T2DM.^4,5^ Adiposity is also a very well-established cause of T2DM.^6^ Whether poor liver function also causes adiposity and thereby contributes to T2DM is unclear. Observationally, poor liver function is associated with obesity,^7,8^ but these studies are open to confounding by lifestyle, including diet^9^ and physical activity,^10^ health status and socioeconomic position (SEP).^11^ As such, whether poor liver function is an additional contributor to the obesity epidemic remains uncertain.

To inform this question when experimental evidence is lacking, we conducted two complimentary analyses with different assumptions and study designs. Observationally, we examined the association of liver function (ALT, alkaline phosphatase (ALP)) with adiposity in young people in a setting with little clear socio-economic patterning of obesity, so as to reduce confounding by poor health and socio-economic position, i.e., in Hong Kong’s “Children of 1997” birth cohort.^12^ We also, for the first time, used an MR study design to assess the effects of liver enzymes on adiposity, which takes advantage of the random allocation of genetic endowment at conception thereby providing randomization analogous to the randomization in randomized controlled trials.^13^ We assessed the associations of genetically predicted liver enzymes (ALT, ALP and gamma glutamyltransferase (GGT))^14^ with adiposity indices, i.e., body mass index (BMI), waist circumference (WC) and waist-hip ratio (WHR), using the Genetic Investigation of ANthropometric Traits (GIANT) consortium.^15-17^ We also considered sex-specific associations, where possible, because sex differences in circulating levels of endogenous sex hormones are associated with both adiposity^18^ and fatty liver.^19^

## Materials and methods

### The “Children of 1997” birth cohort

The “Children of 1997” birth cohort is a population-representative Chinese birth cohort (n=8,327) which included 88% of all births in Hong Kong from 1 April 1997 to 31 May 1997.^20^ The study was initially established to examine the effects of second-hand smoke exposure and breastfeeding on health services utilization to 18 months. Participants were recruited at the first postnatal visit to any of the 49 Maternal and Child Health Centers in Hong Kong, which parents of all newborns were strongly encouraged to attend to obtain free preventive care and vaccinations for their child/children up to 5 years of age. Information, including parental characteristics (maternal age, paternal age, parental smoking and parental education) and infant characteristics (birth weight, gestational age and sex) were obtained from a self-administered questionnaire in Chinese at recruitment and subsequent routine visits. Parental occupation, type of housing and income were also recorded. In 2007, contact was re-established followed by three postal/telephone questionnaire surveys and a Biobank clinical follow-up at 16-18 years. At the Biobank clinical follow-up, as a compromise between cost and comprehensiveness, liver enzymes were assessed from plasma ALT (IU/L) and plasma ALP (IU/L) analyzed using the Roche Cobas C8000 System, a discrete photometric chemistry analyzer, with International Federation of Clinical Chemistry standardized method with pyridoxal phosphate and substrates of L-alanine and 2-oxoglutarate for ALT, and an optimized substrate concentration and 2-amino-2-methyl-1-propanol as buffer plus the cations magnesium and zinc for ALP. These analyses were conducted at an accredited laboratory serving a teaching hospital in Hong Kong. Height (cm), weight (kg) and waist and hip circumference (cm) were measured using standard protocols.

### Children of 1997

#### Exposure - liver function

Liver function at ∼17.5 years was assessed from plasma ALT (IU/L) and plasma ALP (IU/L).

#### Outcome - Adiposity

Adiposity was assessed from BMI (kg/m^2^), WC (cm) and WHR, which represent different aspects of adiposity. Although these are not completely normally distributed, we present them in natural units for ease of interpretation, given interpretation was similar using a gamma distribution.

### Mendelian randomization

#### Genetic associations with liver enzymes

Single nucleotide polymorphisms (SNPs) associated with plasma log transformed ALT, ALP and GGT at genome-wide significance (p-value<5×10^-8^) adjusted for age and sex were obtained from the largest available genome-wide association study (GWAS) of plasma levels of liver enzymes comprising 61,089 adults (∼86% European, mean age 52.8 years, 50.6% women).^14,21^ For SNPs in linkage disequilibrium (R^2^>0.01), we retained SNPs with the lowest p-value using the *Clumping* function of MR-Base (TwoSampleMR) R package, based on the 1000 Genomes catalog.^22^ Whether any of the selected SNPs was related to adiposity directly rather than through liver enzymes (pleiotropic effects) was assessed from their known phenotypes obtained from comprehensive curated genotype to phenotype cross-references, i.e., Ensembl (http://www.ensembl.org/index.html) and the GWAS Catalog (https://www.ebi.ac.uk/gwas/). We also identified SNPs from highly pleiotropic genes, such as *ABO* and *GCKR*, whose full functionality is not yet clearly understood.

#### Genetic associations with adiposity

Overall genetic associations with BMI (standard deviation (SD) units) were obtained from 2018 GIANT and UK Biobank meta-analysis^16^ (GIANTUKB) (n=681,275), a meta-analysis of the GIANT GWAS Anthropometric 2015 BMI^15^ (mean age 56.0 years, 53.8% women, 95% European) with a newly conducted GWAS of UK Biobank (100% European).^16^ Sex-specific genetic associations with BMI were from GIANT GWAS Anthropometric 2015 BMI^15^ (n=339,224, mean age 56.0 years, 53.8% women, 95% European). Overall and sex-specific genetic associations with WC (SD units) and WHR (SD units) were obtained from the GIANT GWAS Anthropometric 2015 Waist^17^ (n=224,459, mean age 54.5 years, 54.6% women, 63.6% European). The GIANT (GWAS Anthropometric 2015 BMI^16^ and the GWAS Anthropometric 2015 Waist^18^) adjusted for age, age-squared, study-specific covariates in a linear model.^15,17^ The UK Biobank adjusted for age, sex, recruitment centre, genotyping batch and 10 principal components.^16^

### Statistical analysis

In the “Children of 1997” birth cohort, baseline characteristics were compared between cohort participants who were included and excluded using Cohen effect sizes,^23^ which indicates the magnitude of difference between groups independent of sample size. Cohen effect sizes are usually categorized as 0.10 for small, 0.30 for medium and 0.50 for large when considering categorical variables. ^23^ The associations of adiposity indices with potential confounders were assessed using independent t-test or analysis of variance.

We assessed the associations of liver function with adiposity indices adjusted for potential confounders, i.e., household income, highest parental education, type of housing, highest parental occupation, second-hand and maternal smoking and sex, using multivariable linear regression. We also assessed whether associations differed by sex from the relevant interaction term.

In the Mendelian randomization study, we estimated the strength of the genetic instruments from the *F*-statistic.^24^ A higher *F*-statistic indicates lower risk of weak instrument bias.^24^ We aligned SNPs for exposure and outcome on allele and effect allele frequency to ensure all SNPs, in particular palindromic SNPs, were aligned correctly. SNPs that could not be unequivocally aligned were replaced by proxies or dropped. SNPs predicting liver function that were not available for adiposity indices were replaced by highly correlated proxies (R^2^>0.9). Potential proxy SNPs were obtained from the GWAS^14^ and their correlations with other SNPs were obtained using LDlink.^25,26^

We obtained unconfounded estimates of the effects of liver enzymes on adiposity indices overall and by sex by combining SNP-specific Wald estimates (SNP-outcome association divided by SNP-exposure association) using inverse variance weighting (IVW) with random effects for 4+ SNPs, which assumes that balanced pleiotropy, and with fixed effects for 3 SNPs or fewer. We repeated the analysis excluding pleiotropic SNPs that might be associated with the relevant outcome directly rather than via liver enzymes. As a sensitivity analysis, we used a weighted median (WM) and MR-Egger regression. The WM may generate correct estimates when >50% of weight is contributed by valid SNPs.^27^ MR-Egger generates correct estimates even when all the SNPs are invalid instruments as long as the instrument strength independent of direct effect assumption is satisfied.^28^ A non-null intercept from MR-Egger indicates potential directional pleiotropy and invalid IVW estimates.^29^ Heterogeneity was assessed using the *I*^*2*^ statistic.^28^

All statistical analyses were conducted using R version 3.4.2 (R Foundation for Statistical Computing, Vienna, Austria). The R package MendelianRandomization^30^ was used to generate the estimates.

#### Ethics approval

Ethical approval for the “Children of 1997” study was obtained from the University of Hong Kong, Hospital Authority Hong Kong West Cluster Joint Institutional Review Board. Ethical approval from an Institutional Review Board is not required for the MR study since it only uses publicly available summary data.

## Results

### Children of 1997

In the Biobank clinical follow-up, 3,460 adolescents of 6,850 potentially active follow-up participants took part (51% follow-up) of whom, 3,458 had at least one measure of BMI, WC or WHR, as shown in Figure 1. The 4,869 participants without adiposity measures were not different from the included participants in terms of sex, second-hand and maternal smoking exposure and SEP with relatively small Cohen effect sizes (<0.13) (Table A.1). The mean and SD of BMI, WC and WHR were 20.9 kg/m^2^ (SD 3.5 kg/m^2^), 72.3 cm (SD 9.2 cm) and 0.77 (SD 0.06). Boys had higher BMI, WC, and WHR than girls. Maternal smoking was associated with larger BMI and WC. SEP had little association with BMI, WC and WHR (Table 1).

**Table 1.** Baseline characteristics by body mass index (BMI), waist circumference (WC) and waist-hip ratio (WHR) among participants in Hong Kong’s “Children of 1997” birth cohort, Hong Kong, China, 1997 to 2016

| Characteristics | BMI (kg/m<sup>2</sup>) |  |  |  | WC (cm) |  |  |  | WHR |  |  |  |
| --- | --- | --- | --- | --- | --- | --- | --- | --- | --- | --- | --- | --- |
|  | No. | % | Mean (SD) | P-value* | No. | % | Mean (SD) | P-value* | No. | % | Mean (SD) | P-value* |
| BMI | 3457 |  | 20.9 (3.5) | - | - | - | - | - | - | - | - | - |
| WC (cm) | - | - | - | - | 3453 | - | 72.3 (9.2) | - | - | - | - | - |
| WHR | - | - | - | - | - | - | - | - | 3452 | - | 0.77 (0.06) | - |
| Sex | 3457 | - | - | <0.001 | 3453 | - | - | <0.001 | 3452 | - | - | <0.001 |
| Girl | 1716 | 49.6% | 20.7 (3.3) | - | 1713 | 49.6% | 69.7 (7.8) | - | 1712 | 49.6% | 0.75 (0.05) | - |
| Boy | 1741 | 50.4% | 21.2 (3.8) | - | 1740 | 50.4% | 74.9 (9.7) | - | 1740 | 50.4% | 0.79 (0.07) | - |
| Unknown | 0 | - | - | - | 0 | - | - | - | 0 | - | - | - |
| Second-hand and maternal smoking exposure | 3457 | - | - | <0.001 | 3453 | - | - | <0.01 | 3452 | - | - | 0.05 |
| None | 943 | 27.3% | 20.6 (3.1) | - | 943 | 27.3% | 71.6 (8.3) | - | 942 | 27.3% | 0.77 (0.06) | - |
| Prenatal second-hand smoking | 1280 | 37.0% | 20.9 (3.7) | - | 1278 | 37.0% | 72.3 (9.5) | - | 1278 | 37.0% | 0.77 (0.06) | - |
| Postnatal second-hand smoking | 961 | 27.8% | 21.3 (3.8) | - | 959 | 27.8% | 73.0 (9.8) | - | 959 | 27.8% | 0.77 (0.07) | - |
| Maternal smoking | 128 | 3.7% | 21.5 (3.4) | - | 128 | 3.7% | 73.6 (9.4) | - | 128 | 3.7% | 0.78 (0.06) | - |
| Unknown | 145 | 4.2% | 20.6 (3.1) | - | 145 | 4.2% | 71.0 (7.8) | - | 145 | 4.2% | 0.76 (0.06) | - |
| Highest parental education level | 3457 | - | - | 0.71 | 3453 | - | - | 0.39 | 3452 | - | - | 0.29 |
| Grade<=9 | 991 | 28.7% | 21.0 (3.8) | - | 990 | 28.7% | 72.6 (9.8) | - | 990 | 28.7% | 0.77 (0.07) | - |
| Grades 10-11 | 1489 | 43.1% | 20.9 (3.4) | - | 1486 | 43.0% | 72.2 (8.8) | - | 1485 | 43.0% | 0.77 (0.06) | - |
| Grades>=12 | 961 | 27.8% | 20.9 (3.5) | - | 961 | 27.8% | 72.3 (9.3) | - | 961 | 27.8% | 0.77 (0.06) | - |
| Unknown | 16 | 0.5% | 20.8 (2.0) | - | 16 | 0.5% | 69.2 (5.1) | - | 16 | 0.5% | 0.74 (0.04) | - |
| Highest parental occupation | 3457 | - | - | 0.01 | 3453 | - | - | 0.07 | 3452 | - | - | 0.42 |
| I (unskilled) | 99 | 2.9% | 20.7 (3.1) | - | 99 | 2.9% | 71.6 (8.9) | - | 99 | 2.9% | 0.77 (0.06) | - |
| II (semiskilled) | 285 | 8.2% | 21.3 (3.7) | - | 284 | 8.2% | 72.8 (9.3) | - | 284 | 8.2% | 0.77 (0.07) | - |
| III M (semiskilled) | 503 | 14.6% | 20.8 (3.5) | - | 503 | 14.6% | 72.2 (9.4) | - | 503 | 14.6% | 0.77 (0.07) | - |
| III NM (nonmanual skilled) | 879 | 25.4% | 21.0 (3.7) | - | 878 | 25.4% | 72.5 (9.4) | - | 877 | 25.4% | 0.77 (0.06) | - |
| IV (managerial) | 440 | 12.7% | 21.4 (3.8) | - | 440 | 12.7% | 73.3 (10.2) | - | 440 | 12.7% | 0.77 (0.07) | - |
| V (professional) | 797 | 23.1% | 20.7 (3.3) | - | 797 | 23.1% | 71.8 (8.7) | - | 797 | 23.1% | 0.77 (0.06) | - |
| Unknown | 454 | 13.1% | 20.7 (3.4) | - | 452 | 13.1% | 71.6 (8.3) | - | 452 | 13.1% | 0.77 (0.06) | - |
| Household income per head at recruitment | 3457 | - | - | 0.12 | 3453 | - | - | 0.14 | 3452 | - | - | 0.45 |
| First quintile | 572 | 16.5% | 20.8 (3.5) | - | 571 | 16.5% | 71.9 (9.0) | - | 571 | 16.5% | 0.77 (0.07) | - |
| Second quintile | 616 | 17.8% | 21.0 (3.7) | - | 614 | 17.8% | 72.4 (9.9) | - | 614 | 17.8% | 0.77 (0.07) | - |
| Third quintile | 618 | 17.9% | 21.2 (3.7) | - | 617 | 17.9% | 73.2 (9.6) | - | 617 | 17.9% | 0.77 (0.06) | - |
| Fourth quintile | 631 | 18.3% | 20.8 (3.4) | - | 631 | 18.3% | 72.1 (8.8) | - | 631 | 18.3% | 0.77 (0.06) | - |
| Fifth quintile | 646 | 18.7% | 20.8 (3.2) | - | 646 | 18.7% | 72.0 (8.6) | - | 645 | 18.7% | 0.77 (0.07) | - |
| Unknown | 374 | 10.8% | 21.1 (3.7) | - | 374 | 10.8% | 72.2 (9.6) | - | 374 | 10.8% | 0.77 (0.07) | - |
| Type of housing at recruitment | 3457 | - | - | 0.18 | 3453 | - | - | 0.51 | 3452 | - | - | 0.47 |
| Public | 1448 | 41.9% | 21.1 (3.6) | - | 1445 | 41.8% | 72.5 (9.4) | - | 1445 | 41.9% | 0.77 (0.07) | - |
| Subsidized home ownership scheme | 545 | 15.8% | 20.8 (3.6) | - | 544 | 15.8% | 72.0 (9.2) | - | 544 | 15.8% | 0.77 (0.06) | - |
| Private | 1359 | 39.3% | 20.9 (3.4) | - | 1359 | 39.4% | 72.3 (9.1) | - | 1358 | 39.3% | 0.77 (0.06) | - |
| Unknown | 105 | 3.0% | 20.5 (3.0) | - | 105 | 3.0% | 71.2 (7.5) | - | 105 | 3.0% | 0.76 (0.05) | - |

\* Two-side *P*-value from independent t-test or analysis of variance (ANOVA)

**Figure 1.**
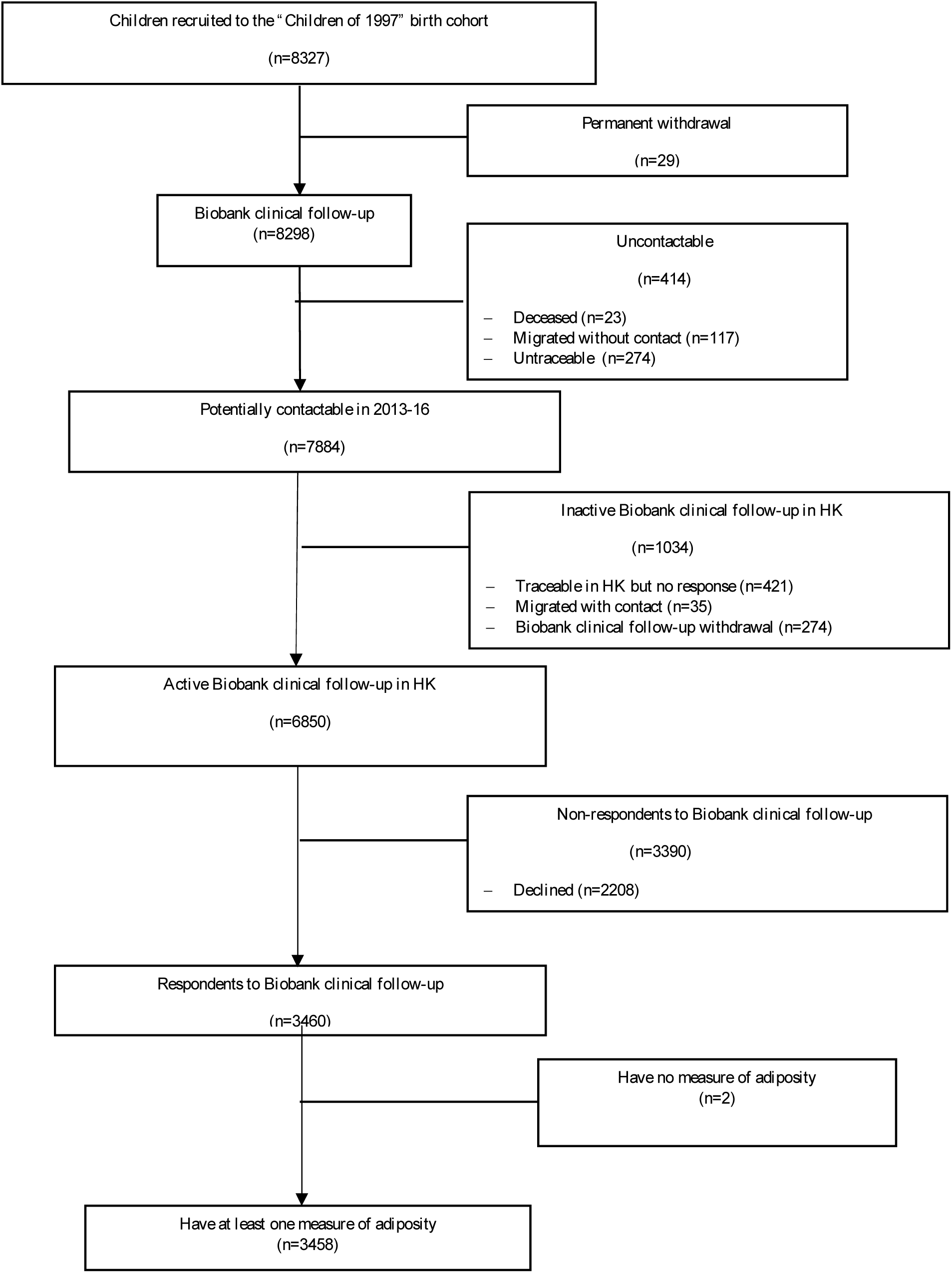
Flowchart of the Hong Kong’s “Children of 1997” birth cohort, Hong Kong, China, 1997 to 2016.

Table 2 shows ALT was positively associated with BMI, WC and WHR adjusted for potential confounders. ALP was negatively associated with BMI, WC and WHR. The associations of ALP with BMI, WC and WHR differed by sex, with the inverse associations only evident in boys.

**Table 2.** Adjusted associations of liver function (alanine aminotransferase (ALT) and alkaline phosphatase (ALP)) with adiposity markers (body mass index (BMI), waist circumference (WC) and waist-hip ratio (WHR)) at ∼17.5 years in the Hong Kong’s “Children of 1997” birth cohort, Hong Kong, China

| Liver enzyme | Outcome | Sex-adjusted |  | Sex interaction<br>p-value | Boys |  | Girls |  |
| --- | --- | --- | --- | --- | --- | --- | --- | --- |
|  |  | Beta | 95% CI |  | Beta | 95% CI | Beta | 95% CI |
| ALT (IU/L) | BMI (kg/m<sup>2</sup>) | 0.10 | 0.09 to 0.11 | 0.19 | 0.10 | 0.09 to 0.12 | 0.09 | 0.07 to 0.11 |
|  | WC (cm) | 0.25 | 0.23 to 0.28 | 0.04 | 0.27 | 0.24 to 0.30 | 0.21 | 0.17 to 0.26 |
|  | WHR | 0.0013 | 0.0012 to 0.0015 | 0.08 | 0.0014 | 0.0012 to 0.0016 | 0.0011 | 0.0008 to 0.0014 |
| ALP (IU/L) | BMI (kg/m<sup>2</sup>) | -0.018 | -0.024 to -0.012 | <0.001 | -0.024 | -0.031 to -0.017 | 0.006 | -0.006 to 0.018 |
|  | WC (cm) | -0.03 | -0.05 to -0.02 | <0.001 | -0.05 | -0.07 to -0.03 | 0.04 | 0.01 to 0.07 |
|  | WHR | -0.0002 | -0.0003 to -0.0001 | 0.002 | -0.0002 | -0.0004 to -0.0001 | 0.0001 | -0.0001 to 0.0003 |

Adjustment: household income, highest parental education, type of housing, highest parental occupation, second-hand and maternal smoking

### Mendelian randomization

#### Genetic variants

In total, 4 SNPs independently predicting ALT, 14 SNPs independently predicting ALP and 26 SNPs independently predicting GGT at genome-wide significance were obtained.^14^ All the palindromic SNPs were aligned based on effect allele frequency (Table A.2), except for rs2073398 (*GGT1, GGTLC2*), predicting GGT, which was replaced by rs5751901 (R^2^=0.95) for GIANTUKB. Rs6834314 (*HSD17B13, MAPK10*) predicting ALT and rs944002 (*EXOC3L4*) predicting GGT which were replaced in the GWAS Anthropometric 2015 Waist by rs13102451 (R^2^=1.00) and rs2297067 (R^2^=0.98). Two SNPs, rs516246 (*FUT2*) and rs8038465 (*CD276*) predicting GGT had rather different allele distributions for GGT and adiposity indices in GIANT (GWAS Anthropometric 2015 BMI^16^ and the GWAS Anthropometric 2015 Waist^17^). They were dropped in a sensitivity analysis separately. No proxy SNP (R^2^>0.9) of rs516246 could be found in GIANTUKB. (Table A.3).

Of the 4 SNPs predicting ALT, rs2954021 (*TRIB1*) predicted both ALT and ALP. Of the 14 SNPs predicting ALP, rs281377 (*FUT2*) is highly associated with resting metabolic rate; rs579459 is located in the *ABO* gene. Of the 26 SNPs predicting GGT, rs516246 (*FUT2*) is associated with obesity-related traits; rs1260326 (*GCKR*) is associated with Crohn’s disease which might be associated with adiposity (Table A.4). The *F* statistics were 15 for ALT, 158 for ALP and 45 for GGT.

#### Genetic associations with BMI, WC and WHR

Genetically instrumented ALT was negatively associations with BMI, with the association more obvious for women. ALT was not associated with WC or WHR. Genetically instrumented ALP and GGT were not clearly associated with BMI, WC or WHR. Overall, there was little statistical evidence of pleiotropy as few MR-Egger intercepts differed from the null. Large *I*^*2*^ were seen for most estimates, heterogeneity was most evident for ALP (Table 3-5).

**Table 3.**
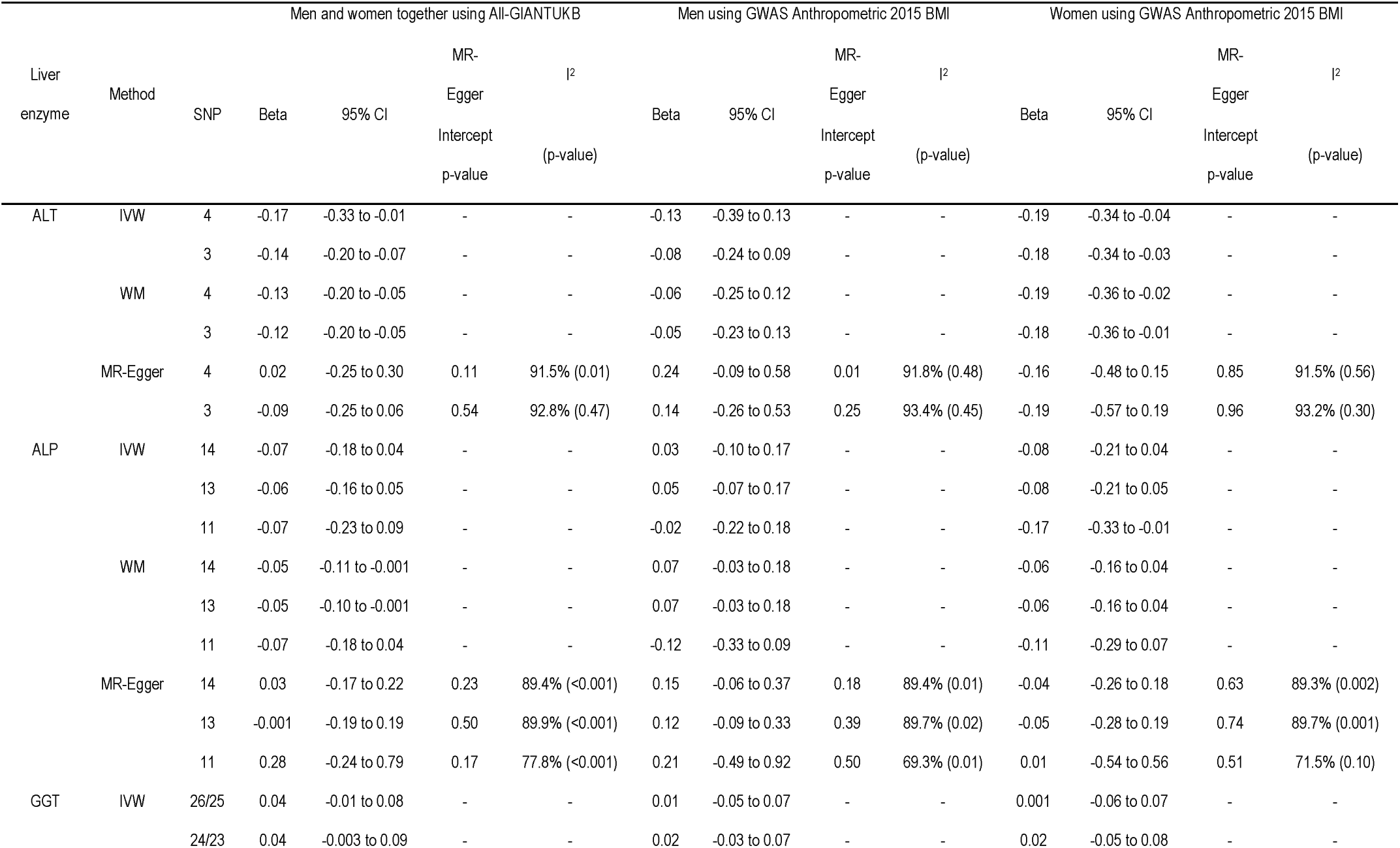
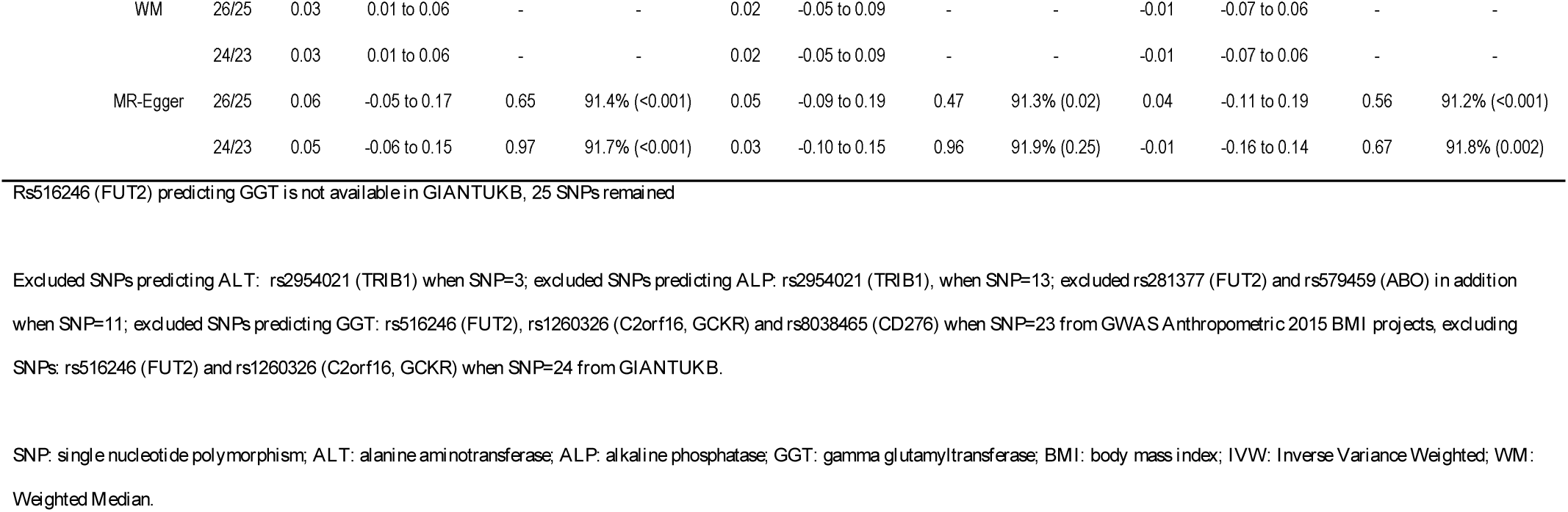
Estimates of the effect of genetically instrumented liver enzymes ALT, ALP and GGT (per 100% change in concentration)^14^ on BMI (standard deviation)^15,16^ using Mendelian randomization with different methodological approaches with and without potentially pleiotropic SNPs

**Table 4.**
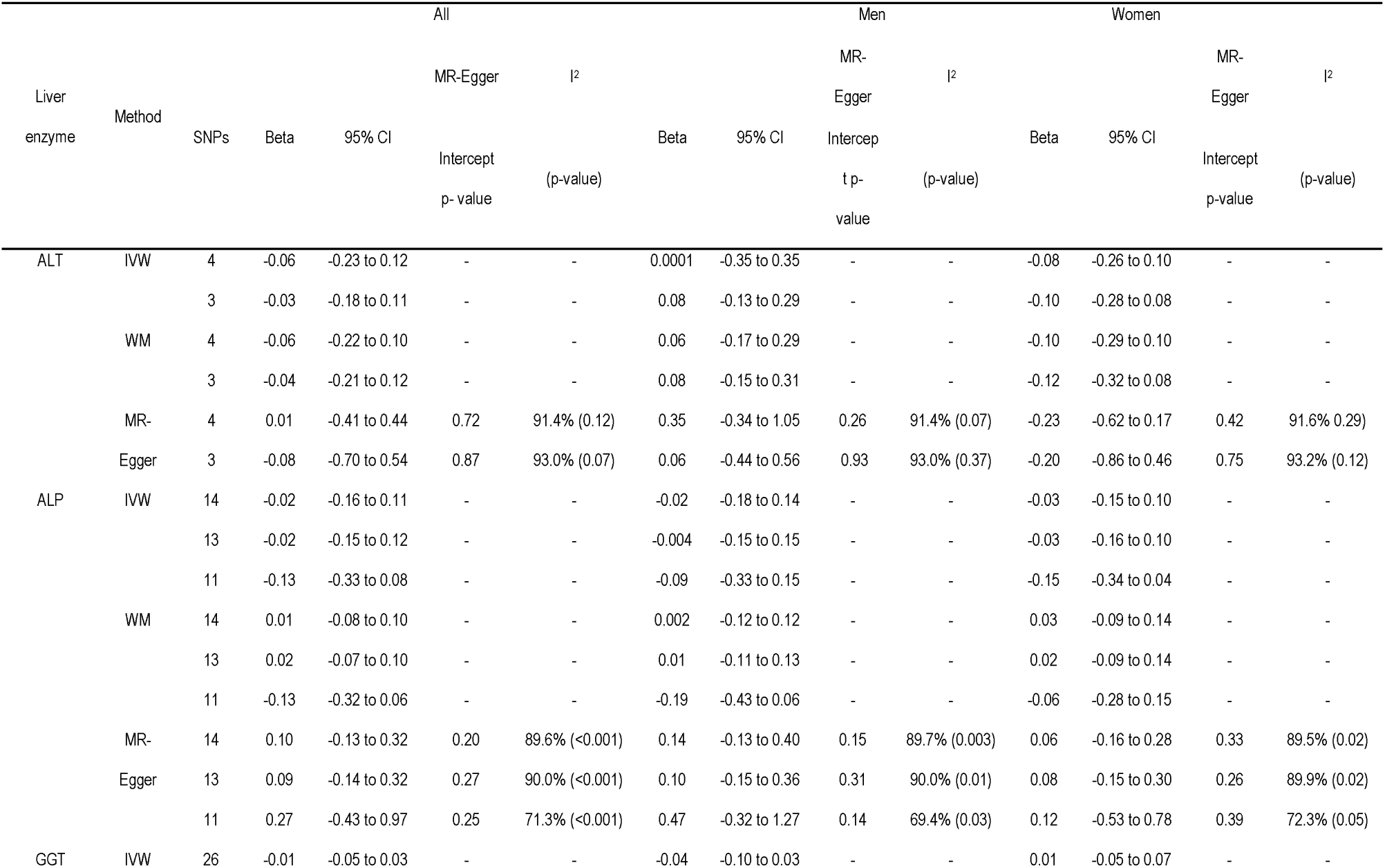
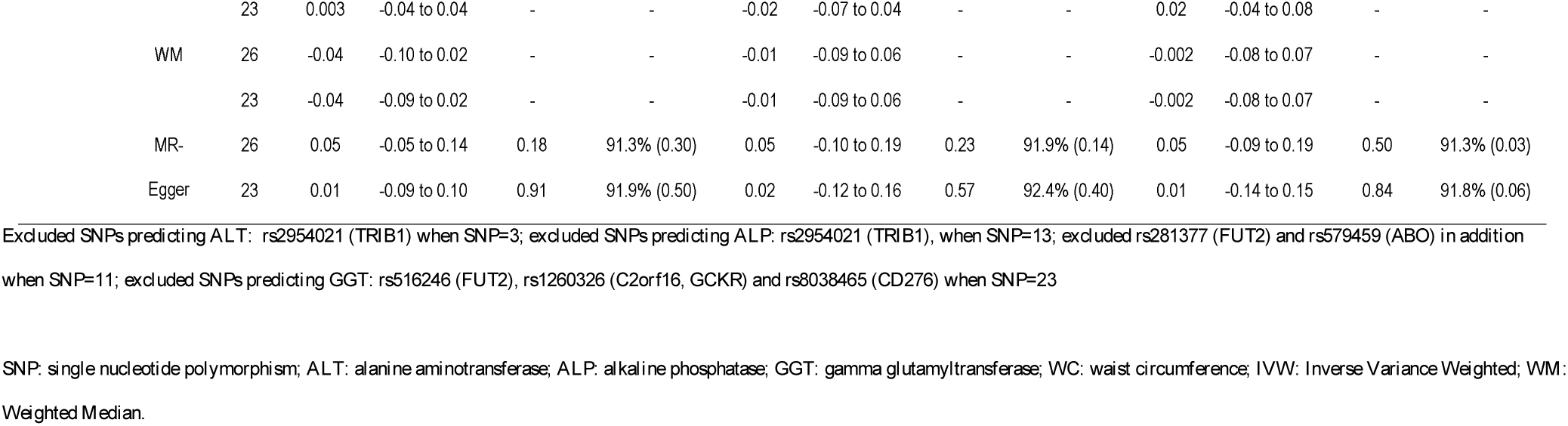
Estimates of the effect of genetically instrumented liver enzymes ALT, ALP and GGT (per 100% change in concentration)^14^ on WC^17^ (standard deviation) using Mendelian randomization with different methodological approaches with and without potentially pleiotropic SNPs

**Table 5.**
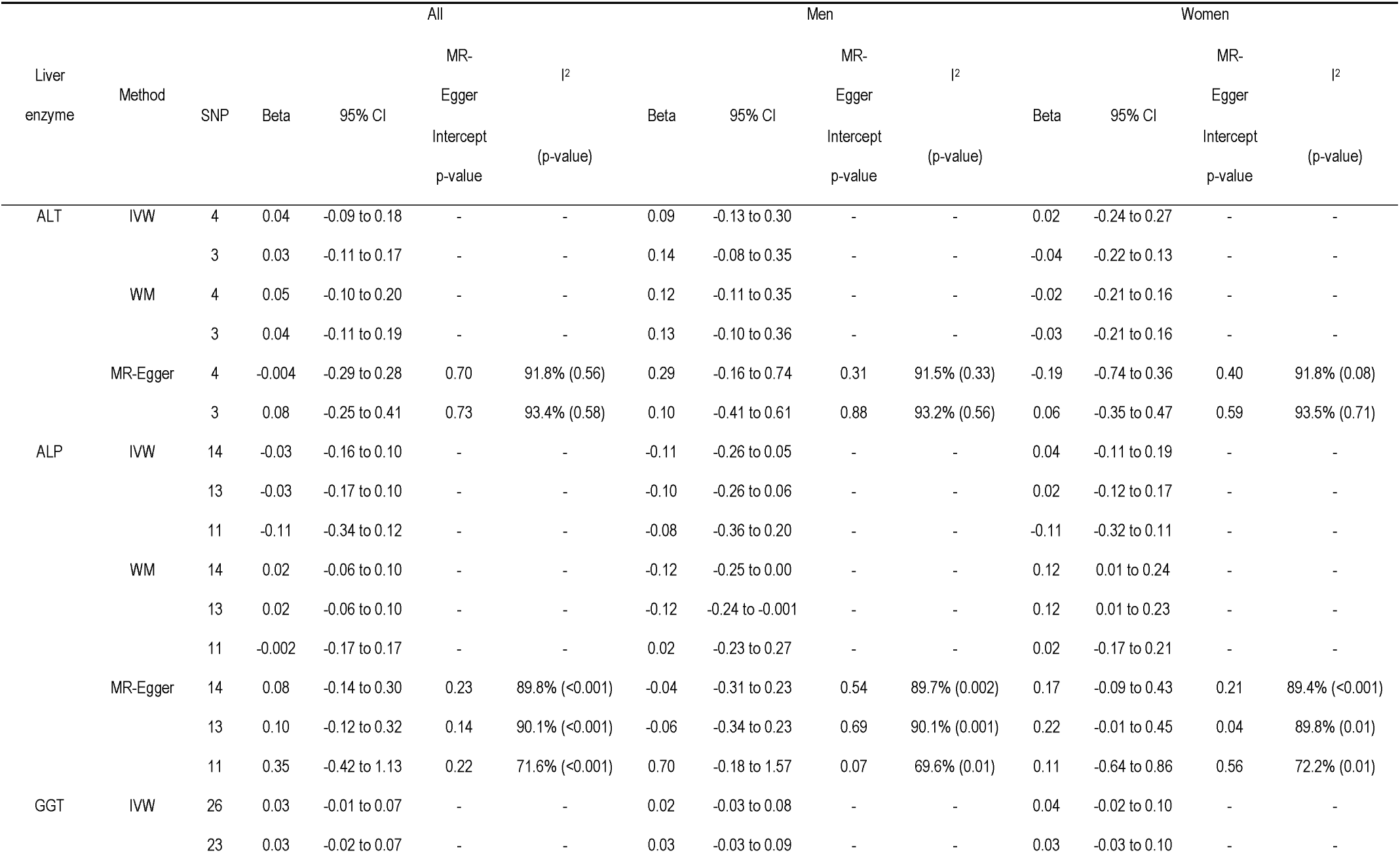
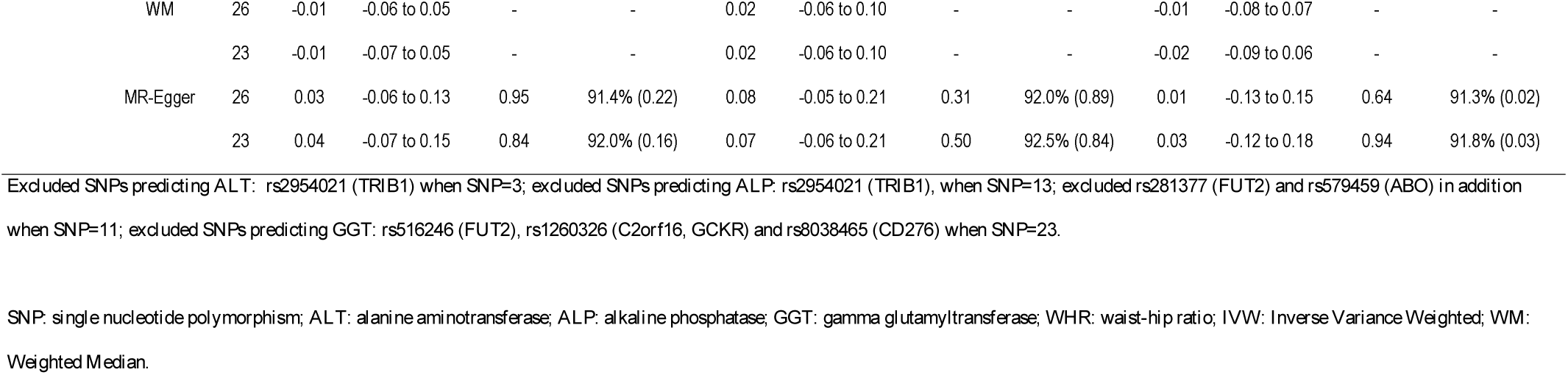
Estimates of the effect of genetically instrumented liver enzymes ALT, ALP and GGT (per 100% change in concentration)^14^ on WHR^17^ (standard deviation) using Mendelian randomization with different methodological approaches with and without potentially pleiotropic SNPs

## Discussion

This novel study used two different approaches, an observational study and an MR study, with different data sources, assumptions, and different unrelated sources of bias to assess the role of liver function in adiposity. We found the clearest evidence for ALT being inversely associated with BMI, perhaps particularly among women.

We used an observational design to assess the association of liver function with adiposity indices in adolescents and an MR design in adults. However, limitations exist. First, observational studies are open to residual confounding by factors such as diet^9^ and physical activity,^10^ which are hard to measure precisely and eliminate. Hong Kong, with a different confounding structure for adiposity, provides a valuable setting in which to triangulate the evidence and to verify observations from Western settings that are potentially confounded.^31^ However, it remains difficult to disentangle correlated factors reliably in an observational study, which might explain discrepancies between observational and MR estimates. It is also possible that associations may vary by history and trajectory of economic development, which has been much more rapid in Hong Kong than in the populations of largely European descent usually included in genetic studies.^32^ Second, ALT was lower than 10 IU/L (n=254) for 7.3% of the participants in “Children of 1997” and was fixed at 5 IU/L, which is unlikely to affect the estimates, because it was only below the limit of detection for a relatively small proportion of observations. Third, follow-up was incomplete, however, no major differences were found between the participants with and without adiposity indices (Table A.1). As such, selection bias from loss-to-follow-up is unlikely be a major concern. Fourth, strong assumptions are required for MR which are also hard to demonstrate empirically, specifically that the genetic instruments, are independent of confounders of the exposure-outcome association and are only associated with the outcome via the exposure. Although we are not certain of the exact function of the SNPs predicting liver enzymes, some of them are mainly expressed in the liver according to the Human Protein Altas (http://www.proteinatlas.org/), making a causal role plausible. Pleiotropic effects are possible, but estimates were similar after excluding potentially pleiotropic SNPs, and MR-Egger did not provide statistical evidence of pleiotropy. Although the *I*^*2*^ was large, it could be driven by the low number of SNPs. Estimates for ALP showed some heterogeneity although the MR-Egger regression did not show directional pleiotropy. Fifth, the overlap of the GWAS for liver enzymes with adiposity indices from GIANT consortium is ∼ 17%, which is unlikely to cause weak instrument bias from any common underlying data structure. Sixth, liver enzymes represent different aspects of liver function: ALT is a marker of hepatocyte integrity, ALP and GGT are markers of cholestasis, but may not completely or only represent liver function.^33^ Seventh, we assessed sex differences on the assumption that genetic predictors of liver function are similar for women and men, which we could not test empirically.^34^

Observationally, the positive associations of ALT with BMI, WC and WHR are consistent with most of the previous observational studies in both adolescents and adults.^35-42^ Observationally, the negative associations of ALP with adiposity are consistent with a previous study among Australian adolescents,^43^ but not with all studies,^44^ although few such studies have been conducted. However, the estimates differed between the observational and MR designs, probably because of the difficulty of distinguishing between correlated measures of liver function and the possibility of confounding. To our knowledge, no previous MR study has assessed the association of liver function with adiposity.

One possible explanation for the finding of ALT potentially reducing BMI is that ALT reduces muscle mass rather than or as well as fat mass, given ALT was inversely associated with BMI but not with WHR or WC. The Korean Sarcopenic Obesity Study, a prospective cohort study, found ALT and NAFLD inversely associated with skeletal muscle mass index.^45^ However, observationally ALT is not consistently associated with sarcopenia,^46^ possibly because of selection bias and confounding in studies of NAFLD patients vulnerable to sarcopenia. However, ALT reducing muscle mass would be consistent with ALT increasing the risk of diabetes,^3^ because low muscle mass is a potential cause of diabetes.^47,48^

## Conclusion

Higher ALT, but not ALP or GGT, possibly reducing specifically BMI, but not WC or WHR suggests that ALT may reduce muscle mass rather than fat mass. Given, MR studies suggest, specifically ALT causes diabetes, whether ALT reduces specifically muscle mass and thereby causes diabetes should be investigated, because it would mean that muscle mass could be an attractive target of intervention to prevent diabetes.

## Acknowledgments

The authors thank colleagues at the Student Health Service and Family Health Service of the Department of Health for their assistance and collaboration. They also thank late Dr. Connie O for coordinating the project and all the fieldwork for the initial study in 1997-1998.

Data on measures of adiposity (2018 GIANT and UK Biobank meta-analysis, GWAS anthropometric 2015 BMI, GWAS anthropometric 2015 waist) have been contributed by the Genetic Investigation of ANthropometric Traits (GIANT) consortium and have been download from https://portals.broadinstitute.org/collaboration/giant/index.php/Main_Page.

## Legends

Table A.1. Baseline characteristics of the participants who were included (n=3,458) and excluded (n=4,869) in the analyses of the Hong Kong’s “Children of 1997” birth cohort, Hong Kong, China, 1997 to 2016

Table A.2. Characteristics of palindromic single nucleotide polymorphisms (SNPs) in the exposure and outcome genome-wide association studies (GWAS)

Table A.3. Characteristics of unequivocally aligned single nucleotide polymorphisms (SNPs) in the exposure and outcome genome-wide association study (GWAS)

Table A.4. Single nucleotide polymorphisms (SNPs) with potential pleiotropic effects other than via the specific liver enzyme from Ensembl and from GWAS Catalog

---

**Table A.1.** Baseline characteristics of the participants who were included (n=3,458) and excluded (n=4,869) in the analyses of the Hong Kong’s “Children of 1997” birth cohort, Hong Kong, China, 1997 to 2016

| Characteristics | Included (n=3,458) | | Excluded (n=4,869) | | Cohen effect size $\Phi$ |
| --- | --- | --- | --- | --- | --- |
|  | n | % | n | % |  |
| Sex | 3458 | - | 4869 | - | 0.08 |
| Female | 1717 | 49.70% | 2196 | 45.10% | - |
| Male | 1741 | 50.30% | 2608 | 53.60% | - |
| Unknown | 0 | - | 65 | 1.30% | - |
| Second-hand and maternal smoking exposure | 3458 | - | 4869 | - | 0.09 |
| None | 943 | 27.30% | 1232 | 25.30% | - |
| Prenatal second-hand smoking | 1280 | 37.00% | 1520 | 31.20% | - |
| Postnatal second-hand smoking | 962 | 27.80% | 1492 | 30.60% | - |
| Maternal smoking | 128 | 3.70% | 275 | 5.60% | - |
| Unknown | 145 | 4.20% | 350 | 7.20% | - |
| Highest parental education levels | 3458 | - | 4869 | - | 0.12 |
| Grade<=9 | 991 | 28.70% | 1476 | 30.30% | - |
| Grades 10-11 | 1490 | 43.10% | 1957 | 40.20% | - |
| Grades>=12 | 961 | 27.80% | 1222 | 25.10% | - |
| Unknown | 16 | 0.50% | 214 | 4.40% | - |
| Highest parental occupation | 3458 | - | 4869 | - | 0.07 |
| I(unskilled) | 99 | 2.90% | 140 | 2.90% | - |
| II(semiskilled) | 285 | 8.20% | 441 | 9.10% | - |
| III M (semiskilled) | 504 | 14.60% | 711 | 14.60% | - |
| III NM (nonmanual skilled) | 879 | 25.40% | 1167 | 24.00% | - |
| IV (managerial) | 440 | 12.70% | 682 | 14.00% | - |
| V(professional) | 797 | 23.00% | 917 | 18.80% | - |
| Unknown | 454 | 13.10% | 811 | 16.70% | - |
| Household income per head at recruitment | 3458 | - | 4869 | - | 0.07 |
| First quintile | 573 | 16.60% | 878 | 18.00% | - |
| Second quintile | 616 | 17.80% | 868 | 17.80% | - |
| Third quintile | 618 | 17.90% | 811 | 16.70% | - |
| Fourth quintile | 631 | 18.20% | 788 | 16.20% | - |
| Fifth quintile | 646 | 18.70% | 794 | 16.30% | - |
| Unknown | 374 | 10.80% | 730 | 15.00% | - |
| Type of housing at recruitment | 3458 | - | 4869 | - | 0.08 |
| Public | 1448 | 41.90% | 2128 | 43.70% | - |
| Subsidized home ownership scheme | 545 | 15.80% | 580 | 11.90% | - |
| Private | 1360 | 39.30% | 1885 | 38.70% | - |
| Unknown | 105 | 3.00% | 276 | 5.70% | - |

Φ Cohen effect sizes are usually categorized into 3 levels, Chi-square tests for categorical variables: 0.10 for small, 0.30 for medium, 0.50 for large. For categorical variables, Cohen's w effect size is calculated as $w = \sqrt{\sum (p_0 - p_1)^2 / p_0}$ , where p0 is the proportion in given by the null hypothesis and p1 is the proportion given the alternative hypothesis; $w = \sqrt{\chi^2 / N}$ where N is the total count of the included and excluded participants.<sup>23</sup>

**Table A.2.** Characteristics of palindromic single nucleotide polymorphisms (SNPs) in the exposure^14^ and outcome genome-wide association studies (GWAS)^15–17^

| Phenotype | SNP | Effect Allele | Other Allele | EAF | EAF_GIANT<sup>15,17</sup> | EAF_GIANTUKB<sup>16</sup> |
| --- | --- | --- | --- | --- | --- | --- |
| ALT | rs10883437 | T | A | 0.64 | 0.62 | 0.61 |
| ALT | rs738409 | G | C | 0.23 | 0.23 | 0.23 |
| ALP | rs10819937 | C | G | 0.17 | 0.14 | 0.19 |
| ALP | rs6984305 | A | T | 0.11 | 0.11 | 0.11 |
| ALP | rs7186908 | C | G | 0.24 | 0.18 | 0.2 |
| GGT | rs2073398 | G | C | 0.34 | 0.39 | NA |
| GGT | rs754466 | T | A | 0.24 | 0.29 | 0.26 |
| GGT | rs9913711 | C | G | 0.65 | 0.68 | 0.66 |

ALT: alanine aminotransferase; ALP: alkaline phosphatase; GGT: gamma glutamyltransferase
EAF: effect allele frequency obtained from Chambers et al., 2011<sup>14</sup>
EAF\_GIANT: effect allele frequency obtained from GWAS Anthropometric 2015 BMI<sup>15</sup> and GWAS Anthropometric 2015 Waist<sup>17</sup>
EAF\_GIANTUKB: effect allele frequency obtained from the 2018 GIANT and UK Biobank meta-analysis<sup>16</sup>
NA: no available data in specific dataset

**Table A.3.**
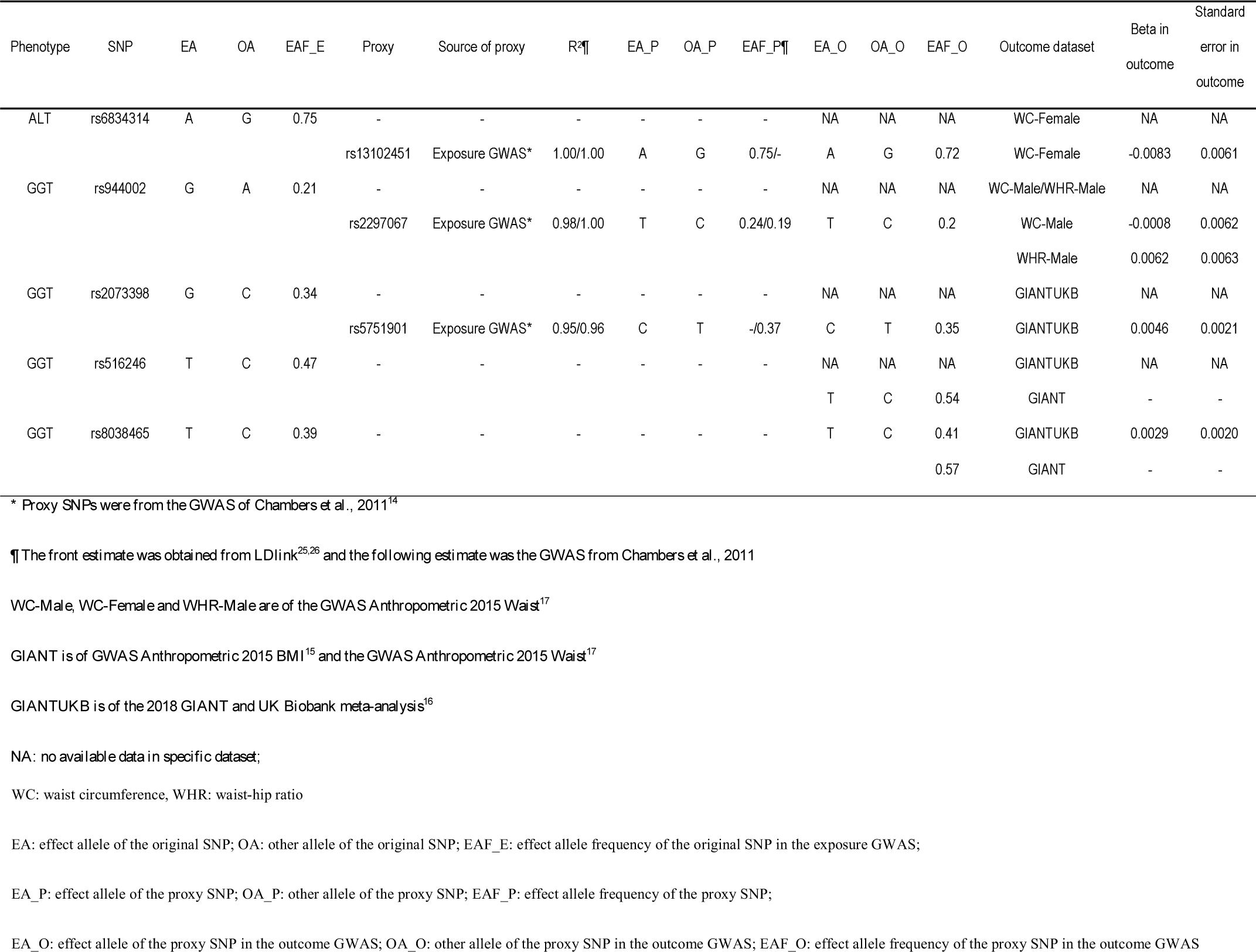
Characteristics of unequivocally aligned single nucleotide polymorphisms (SNPs) in the exposure^14^ and outcome genome-wide association study (GWAS)^15–17^

**Table A.4.**
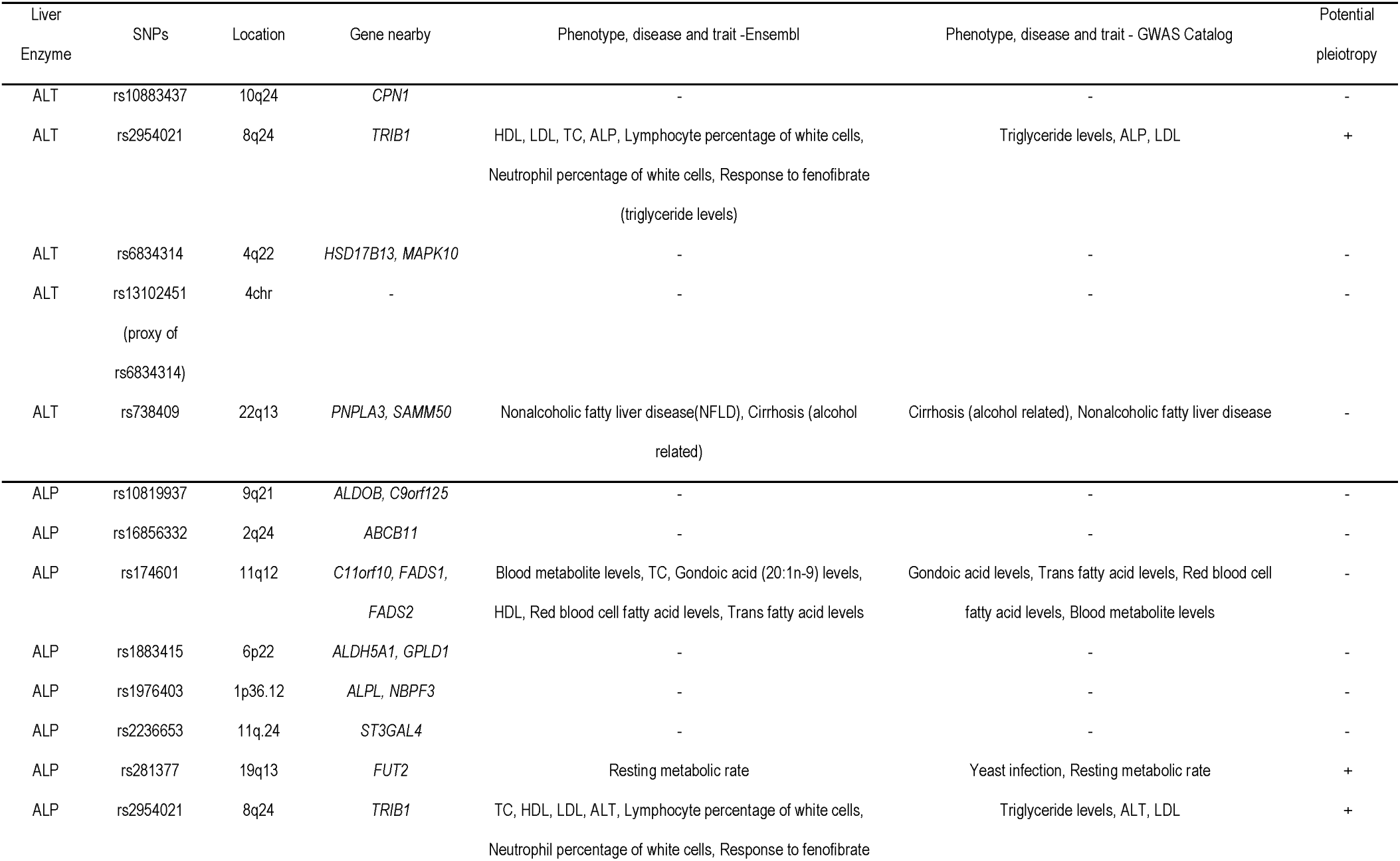
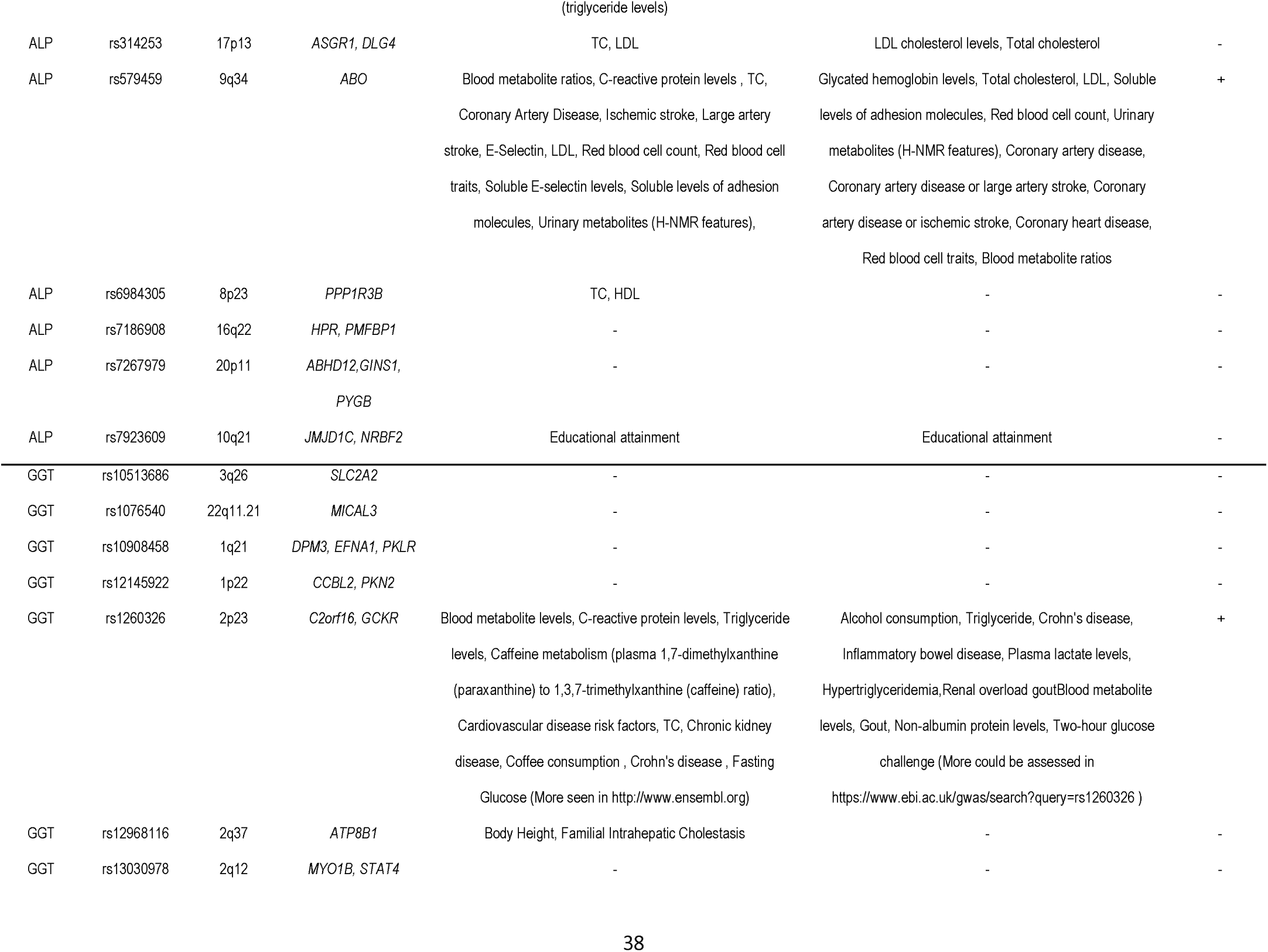
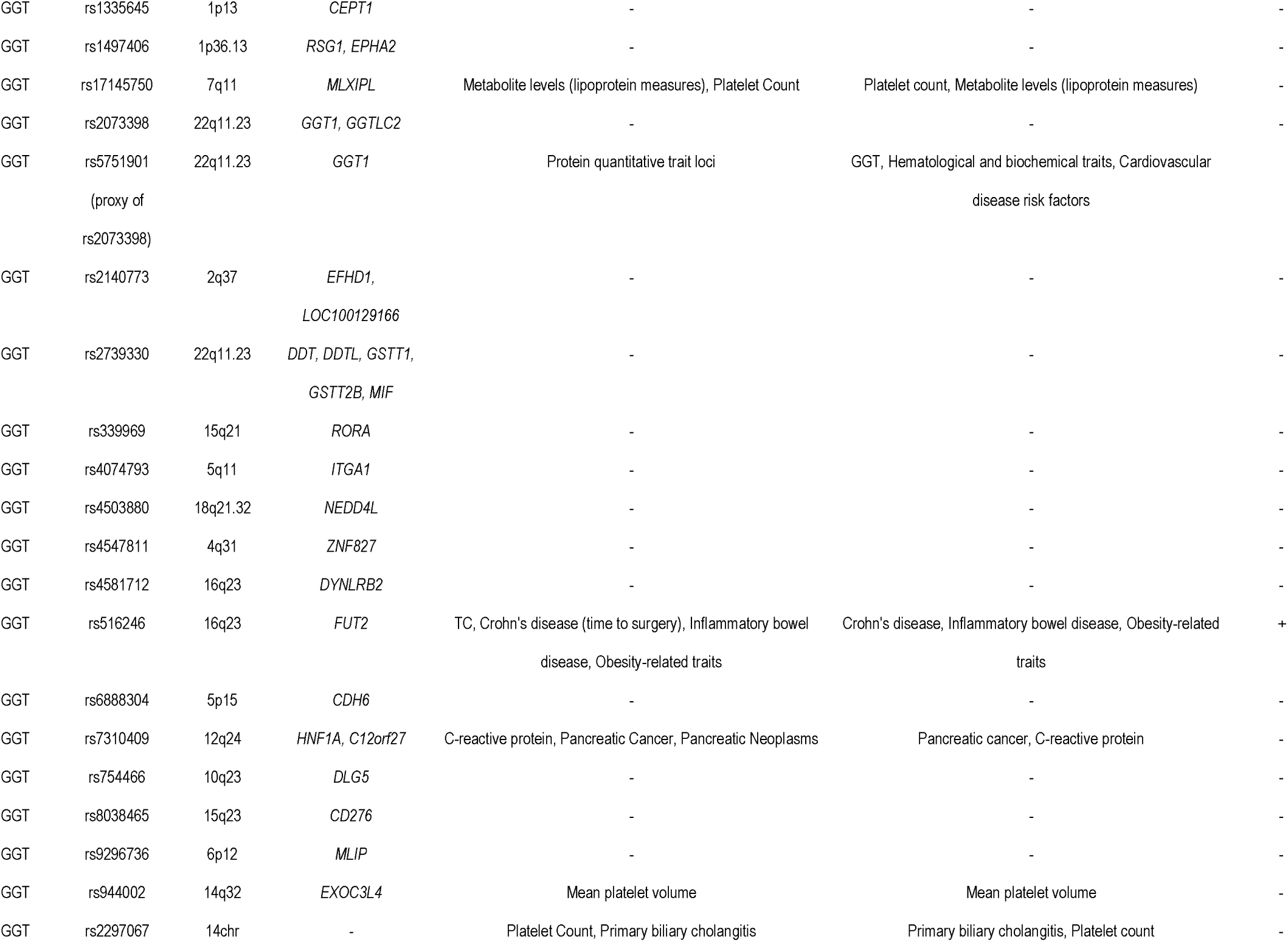
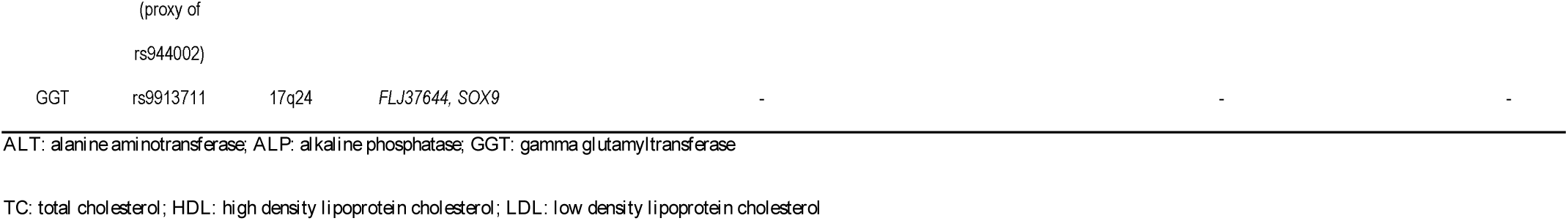
Single nucleotide polymorphisms (SNPs) with potential pleiotropic effects other than via the specific liver enzyme from Ensembl and from GWAS Catalog

